# Individual-specific memory reinstatement patterns within human face-selective cortex

**DOI:** 10.1101/2023.08.06.552130

**Authors:** Yvonne Y. Chen, Aruni Areti, Daniel Yoshor, Brett L. Foster

## Abstract

Humans have the remarkable ability to vividly retrieve sensory details of past events. According to the theory of sensory reinstatement, during remembering, brain regions involved in the sensory processing of prior events are reactivated to support this perception of the past. Recently, several studies have emphasized potential transformations in the spatial organization of reinstated activity patterns. In particular, studies of scene stimuli suggest a clear anterior shift in the location of retrieval activations compared with those during perception. However, it is not clear that such transformations occur universally, with evidence lacking for other important stimulus categories, particularly faces. Critical to addressing these questions, and to studies of reinstatement more broadly, is the growing importance of considering meaningful variations in the organization of sensory systems across individuals. Therefore, we conducted a multi-session neuroimaging study to first carefully map individual participants face-selective regions within ventral temporal cortex (VTC), followed by a second session to examine the correspondence of activity patterns during face memory encoding and retrieval. Our results showed distinct configurations of face-selective regions within the VTC across individuals. While a significant degree of overlap was observed between face perception and memory encoding, memory retrieval engagement exhibited a more selective and constricted reinstatement pattern within these regions. Importantly, these activity patterns were consistently tied to individual-specific neural substrates, but did not show any consistent direction of spatial transformation (e.g., anteriorization). To provide further insight to these findings, we also report on unique human intracranial recordings from VTC under the same experimental conditions. Our findings highlight the importance of considering individual variations in functional neuroanatomy in the context of assessing the nature of cortical reinstatement. Consideration of such factors will be important for establishing general principles shaping the neural transformations that occur from perception to memory.

## Introduction

During episodic memory retrieval, rich sensory details of prior events and experiences can be consciously remembered to support behavior. A leading theory for this process posits that accessing sensory details of past events occurs via the “reinstatement” of prior cortical activity patterns that were present during those events (i.e., encoding) (Danker & Anderson, 2010; Tulving & Thomson, 1973). This sensory reinstatement of prior encoding patterns is thought to result from hippocampally mediated cortical reactivation (Bosch et al., 2014; McClelland et al., 1995; Staresina & Wimber, 2019; Xue, 2018). A substantial literature has reported how encoding activity patterns in sensory cortex are reinstated during retrieval in a stimulus-specific fashion, which is strongly linked to memory behavior, whereby the degree of similarity between retrieval and encoding predicts greater memory strength and accuracy (Favila et al., 2018; Gordon et al., 2014; Kuhl & Chun, 2014; Polyn et al., 2005; Ritchey et al., 2013; St-Laurent et al., 2015; Wang et al., 2022). Subsequently, a commonly held view suggests that the fidelity of encoding activity pattern reinstatement during retrieval is associated with the accuracy of memory contents. A natural extension of this view would be to consider the case of “perfect” reinstatement, where an identical pattern of encoding activity is reinstated during retrieval. This scenario would result in a vivid hallucination, making memory contents indistinguishable from perceptual encoding. However, human memory retrieval is not a perfect recapitulation of prior perceptual experiences, but rather a transformed and reconstructed form of prior encoding (Bartlett & Burt, 1932; Hemmer & Steyvers, 2009; Roediger, 2001; Squire, 2004). Therefore, in more recent years, a growing literature has focused on identifying what principles and mechanisms support these transformations between perception and memory for the same stimuli (Favila et al., 2020; Xue, 2022).

Studies of cortical reinstatement during episodic retrieval have heavily focused on memory-driven reactivation of visual category domains, particularly face and place stimuli, given their well-known and distinct selectivity within the ventral temporal cortex (VTC) (Danker et al., 2011; Gordon et al., 2014; Kuhl et al., 2012; Kuhl & Chun, 2014; Polyn et al., 2005; Prince et al., 2009). By providing a clear and identifiable encoding substrate, the reinstatement of sensory categories in VTC can be carefully examined to test the nature of cortical reinstatement and transformations. Most recently, growing evidence suggests a potential transformation of cortical responses between perception and memory, whereby a topographical anterior shift in the locus of retrieval activations occurs relative to the encoding substrate. This “anterior shift” has precedents in the literature across different stimulus domains (Beauchamp et al., 1999; Chao & Martin, 1999; Hsu et al., 2011; Rugg & Thompson-Schill, 2013; Simmons et al., 2007), however, most recently supporting evidence has come from studies of place/scene perception and memory (Bainbridge et al., 2021; Baldassano et al., 2016; Fairhall et al., 2014; Silson et al., 2016, 2019; Srokova et al., 2022; Steel et al., 2021). At present, it is unclear if this mnemonic anteriorization for retrieval is also consistently observed for face stimuli and reflects a common feature of reinstatement (Gordon et al., 2014; Lee & Kuhl, 2016; Mundy et al., 2013; Schultz et al., 2022; Taylor et al., 2007). As reinstatement is proposed to recapitulate encoding patterns, the identification and quantification of any transformation requires careful delineation of the encoding substrate of each individual. In addressing this question, it is important to consider advances made in identifying individual variations in the organization of visual category-responsive regions throughout VTC.

Historically, response selectivity for face stimuli was associated with a localized aspect of the fusiform gyrus within the human VTC, defining the fusiform face area (FFA) (Kanwisher, 2010, 2017; Kanwisher et al., 1997; Kanwisher & Yovel, 2006). Subsequent work in the human and non-human primate has shown there to be a more complex network of face-selective regions (patches) throughout the ventral visual stream (Chang & Tsao, 2017; Chen et al., 2023; Hesse & Tsao, 2020; Pinsk et al., 2009; Tsao et al., 2006, 2008; Weiner & Grill-Spector, 2010, 2012, 2013). Importantly, this progress includes evidence of classic FFA being comprised of two distinct face-selective clusters in the fusiform gyrus: mFus (medial fusiform, FFA2) and pFus (posterior fusiform, FFA1) (Chen et al., 2023; Pinsk et al., 2009; Weiner & Grill-Spector, 2010, 2012, 2013), often observed together with the inferior occipital gyrus (IOG, or “occipital face area”: OFA) (Gauthier et al., 2000; Pitcher et al., 2011; Rossion et al., 2018). Most recently, large-scale examination has shown systematic variation in the number and configuration of face-selective regions, suggesting three putative “types” of face-selective organization (Chen et al., 2023). As the encoding substrate, it is critical to account for such individual variation when identifying and comparing encoding-retrieval activity patterns. Standard group-level analysis approaches may fail to fully capture these individual differences in encoding-retrieval similarities, potentially distorting the detection of topographic activity transformations. Therefore, in the present study, we performed a multi-session functional magnetic resonance imaging (fMRI) experiment to examine the organization of face selectivity in the human ventral stream during face perception, memory encoding, and retrieval (Figure 1a,b). This approach was designed to study these responses with regard to individuals, allowing for a more careful assessment of reinstatement patterns and their putative transformations, such as anteriorization, from encoding.

**Figure 1.**
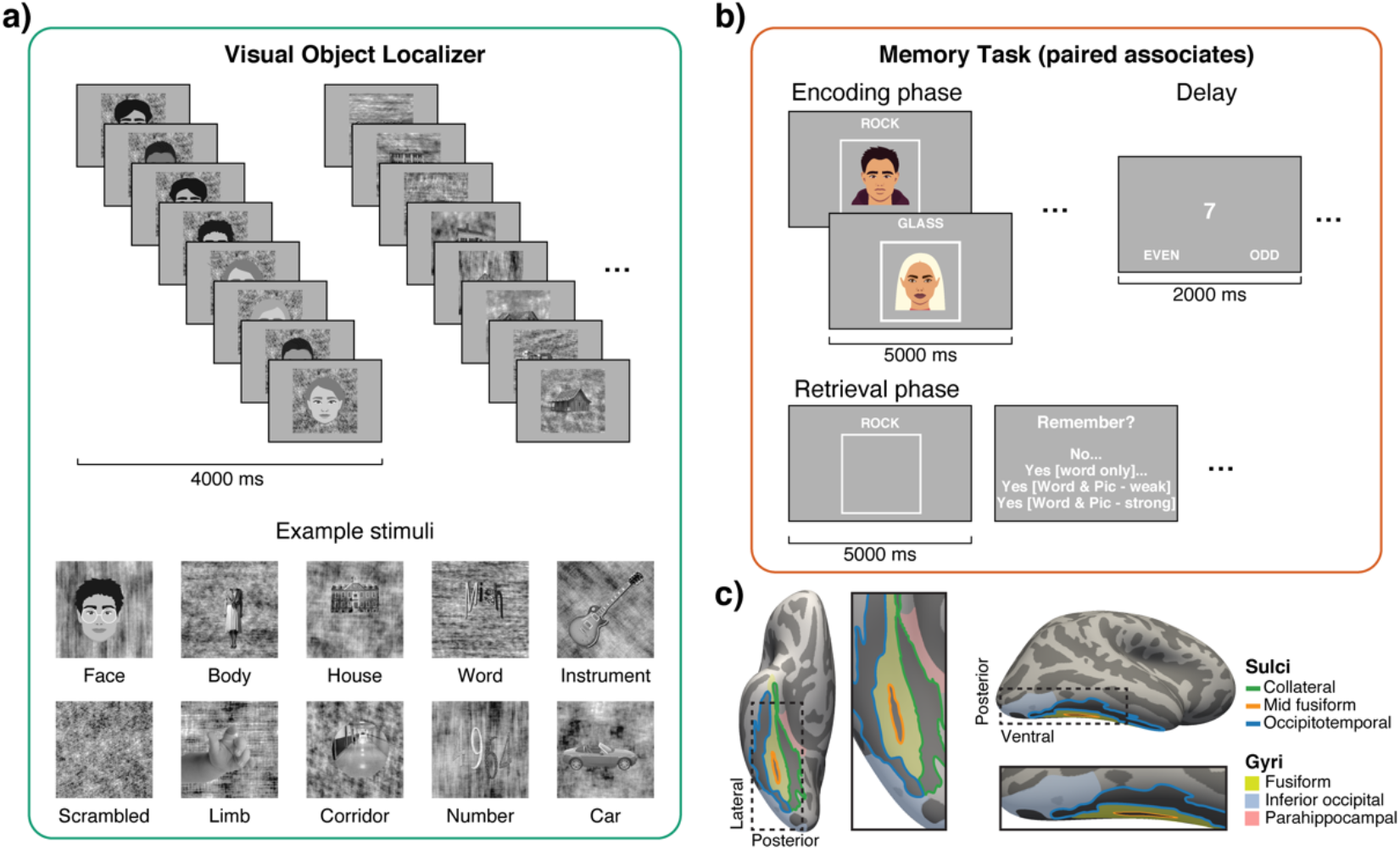
Experimental tasks and region of interest. **a)** Visual object localizer task for identifying face-selective regions. During each trial, eight images from a given visual category were presented as a mini block at the rate of 500 ms per image for a total of 4,000 ms. Participants had to provide a button press to indicate any 1-back stimulus repetition. Example stimuli (bottom) from each of the 10 visual categories are shown. **b)** Memory task experimental procedure. During the encoding phase, word-image pairs were presented for 5,000 ms. During the delay period, between the encoding and retrieval phases, participants were asked to make an odd or even judgment on presented single-digit numbers. During the retrieval phase, cue words (old/studied and new/unstudied) were presented for 5,000 ms followed by a memory strength judgment. Participants were required to encode word-face associations and to retrieve the associated face image from previously studied word cues. **c)** Example right hemisphere anatomy of the human ventral temporal cortex (inflated), highlighting key sulci and gyri.

Through this approach, we identified the organization of face-selective cortex within individuals and found clear evidence for three “types” of configuration at rates consistent with the population data (Chen et al., 2023). Next, we examined how this putative face processing substrate is engaged during memory encoding and retrieval. Importantly, no face stimuli were presented during retrieval, such that any category selectivity reflected retrieval/reinstatement-related processes. We observed that memory encoding of face stimuli recapitulated much of the identified face-selective regions. During retrieval, we observed consistent overlap of encoding regions, but with a reduced magnitude and spatial extent of response - consistent with mnemonically vs. sensory-driven activity. Importantly, we did not find any evidence, beyond contraction, of any consistent topographic transformations, including anteriorization, of retrieval activity patterns. Finally, in consideration of the neurophysiological basis of our observed fMRI responses, and also in regard to understanding the temporal domains of perception-memory transformations, we report on unique human intracranial electroencephalogram (iEEG) data where we examined VTC topography during the same experimental conditions. Together these findings support the view that, for face category stimuli, memory reinstatement recapitulates individual specific encoding substrates, but does not appear to reflect any consistent directed topographic transformation.

## Results

### Task Performance

Participants underwent two sessions of fMRI scanning. In the first session, visual object localizer, participants viewed images from ten visual categories and were instructed to press a button when the same image was presented back-to-back (1-back task, as shown in Figure 1a). Participants performed well on the task, with a mean accuracy of 93.95% (s.d.: ±10.41).

In the second session, the paired associates memory task, participants were presented with word-face pairs for encoding. Later, they were given cue words and asked to retrieve the associated faces. The cue words consisted of words that were paired with faces during encoding (referred to as “old” cues) and words that were not (referred to as “new” cues). Participants were instructed to make one of four responses to each cue word: 1) “No” - if they did not remember the cue word from the encoding phase, 2) “Yes [word only]” - if they remembered the cue word but not the associated face, 3) “Yes [word & pic - weak]” - if they remembered the cue word and had some details about the paired face, 4) “Yes [word & pic - strong]” - if they remembered the cue word and retrieved vivid details about the paired face (Figure 1b). Memory accuracy was assessed based on participants’ responses. Hits referred to correctly identifying the old cue words by responding “yes” (responses 2-4), while correct rejections referred to correctly identifying new cue words by responding “no.” A corrected memory performance score (*d’*) was also calculated. Overall, participants exhibited high memory performance, with a hit rate of 85.2% (s.d.: ±14.6), a correct rejection rate of 90.1% (s.d.: ±12.7), and a *d’* value of 2.8 (s.d.: ±0.87).

### Face perception and memory encoding in ventral temporal cortex face-selective regions

Growing evidence has shown that face-selectivity within the VTC is not limited to a single focal region but encompasses several distinct face-selective areas or “patches” in the human and non-human primate (Chang & Tsao, 2017; Chen et al., 2023; Hesse & Tsao, 2020; Pinsk et al., 2009; Tsao et al., 2006, 2008; Weiner & Grill-Spector, 2010, 2012, 2013). Indeed, a recent large-scale study involving 1,000 participants revealed systematic differences in the configuration of face-selective regions within human VTC, such that individual activation maps did not resemble the averaged group map (Chen et al., 2023). Whereby, this study identified three types of face-selectivity configurations: i) “separate” type with a sizable cortical gap between mFus and pFus; ii) “continuous” type with little to no cortical gap between these regions; iii) “single” type containing only mFus or pFus defined by anatomical landmarks.

In order to examine the nature of functional overlap between memory encoding and retrieval, we followed the common approach of leveraging visual category stimuli, specifically face stimuli, given their behavioral relevance to episodic memory and well know functional organization in VTC, as noted above. To address this question at an individual level, we first had a separate functional visual object localizer scan (session 1) to carefully and precisely identify face-selective cortices and their spatial configuration for each participant. In doing so, we obtained an individual-specific template of “face-selective” regions for subsequent encoding and retrieval activity comparisons.

First, we identified face-selective regions within each participant, specifically the three face-selective areas: mFus, pFus, and IOG. To identify face-selective cortex, we used a general linear model (GLM) to identify regions where responses to face stimuli were greater than nine other visual categories shown (bodies, cars, corridors, instruments, limbs, houses, letters, numbers, and phase-scrambled noise). Face-selective clusters were identified based on a common t-statistic threshold across participants (t>3, vertex level). Within each subject, face-selective functional clusters were labeled based on their anatomical location and spatial extent, and consequently, the ‘type’ of face-selectivity configuration (separate, continuous, and single) established by Chen et al. (2023) was assigned. Figure 2a shows the organization of these three face-selective regions on the inflated cortical surfaces of three example participants. We reliably identified three face-selective regions (mFus, pFus, and IOG) in the majority of participants in both left and right hemispheres (Figure 2e). Additionally, we observed three types of face-selectivity configurations: the separate type (73.53%, Figure 2a top), the continuous type (5.88%, Figure 2a middle), and the single type (14.71%, Figure 2a bottom). Within our sample (n=16), the proportions of spatial configuration types closely matched those previously reported in a large cohort (Chen et al., 2023).

**Figure 2.**
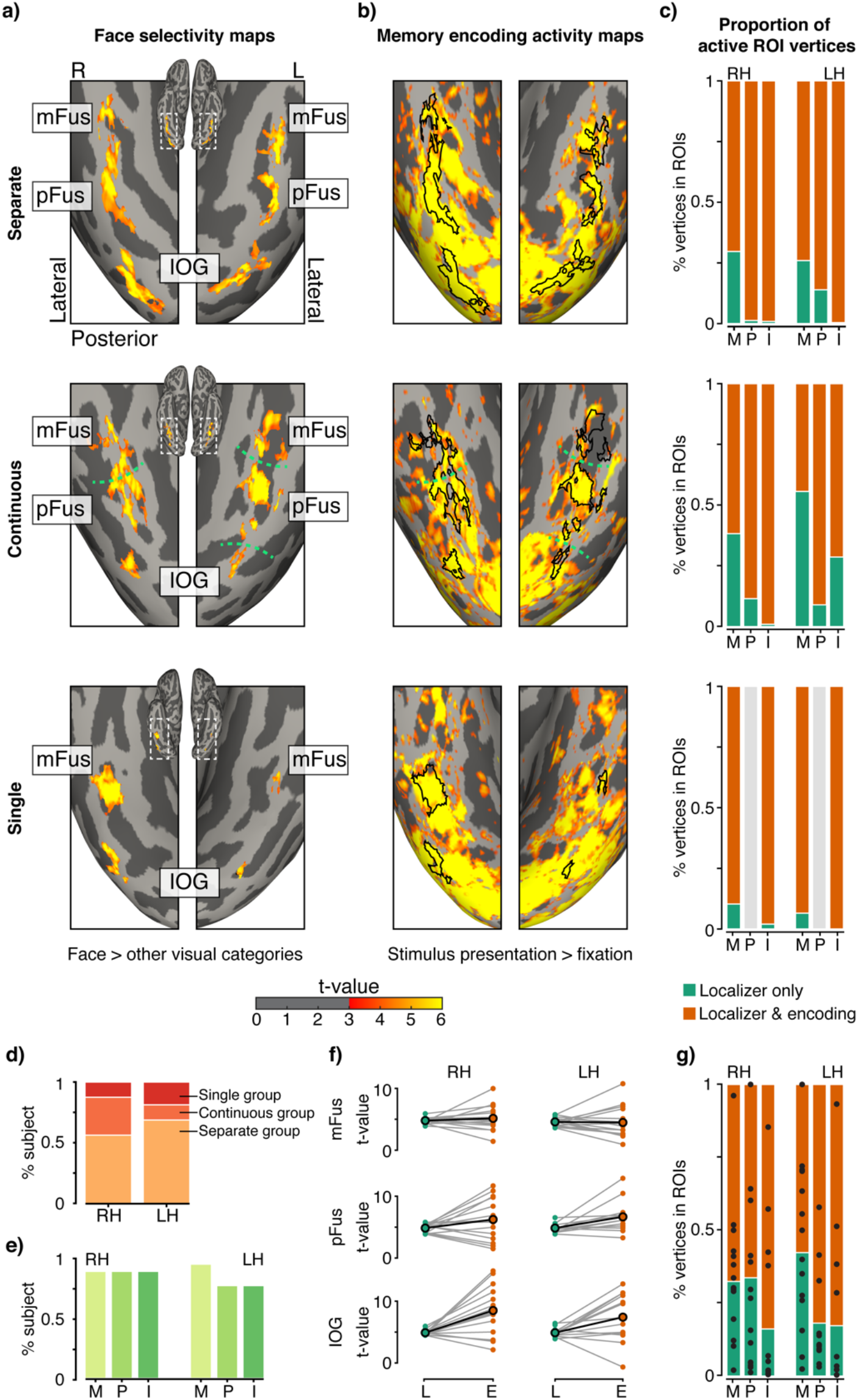
Face-selectivity and memory encoding activity in ventral temporal cortex. **a)** Example participants showing face selectivity in ventral temporal cortex (GLM contrast map: face > all other visual categories). Three types of face-selective-region spatial configurations were observed: separate type (top) with discrete middle Fusiform (mFus), posterior Fusiform (pFus), and inferior occipital gyrus (IOG) face-selective regions; continuous type (middle) with a continuous mFus and pFus region and IOG; and single type (bottom) with only mFus or pFus region and IOG. A common threshold (t >3, vertex level) was used to define all regions across all participants. Color maps illustrate the t-statistics of activation for the given contrast. **b)** Memory encoding activities (GLM contrast map: stimulus presentation > fixation) for the same separate, continuous, and single type example participants in (a). Black outlines in (b) denote the boundaries of face-selective regions from (a). **c)** Proportions of vertices within each face-selective region (mFus(M), pFus(P) and IOG(I)) that were active during localizer only (green) or both localizer and memory encoding (orange) for the same example participants in (a; b). **d)** Percentage of participants with each type of spatial configuration of face-selective regions for left and right hemispheres. **e)** Percentage of participants with face-selective regions identified in left and right hemispheres. **f)** Mean t-value of GLM contrasts (Localizer (L): Face > other visual categories; Memory Encoding (E): Stimulus presentation > inter-trial fixation) are shown for all participants and across three face-selective regions in both left and right hemispheres. **g)** Group averaged proportions of vertices in each face-selective region that were active during localizer only (green) or both localizer and memory encoding (orange). Each black dot denotes the proportion of vertices active during both localizer and memory encoding for a given participant.

Having identified face-selective areas in each participant, we next proceeded to examine the extent to which this neural substrate was engaged during the encoding of faces. During the encoding phase of a separate memory task (session 2), participants were presented with word-face pairs (Figure 1b), with the images of faces expected to elicit responses in the identified face-selective regions. Using a GLM contrast of encoding stimulus presentation > fixation during the memory task, we identified memory encoding activation maps that revealed the brain areas involved in encoding and processing of word-face pairs. Figure 2b displays the associated memory encoding activity maps for the three example configuration types, with putative face-selective regions previously identified outlined in black. Encoding activation maps were generated using a common t-statistic threshold across participants (t > 3, vertex level). Of the three sample participants, large portions of VTC showed strong encoding activity (yellow-red colormap).

To further quantify the spatial overlap of responses, we examined the degree of activity overlap and the activation magnitude between face perception and memory encoding. First, using face-selective regions as anatomical masks, we identified vertices as active if their memory encoding t-statistic was greater than 3. Figure 2c shows the proportions of active vertices during both face perception and memory encoding, as well as face perception alone, for the three example configuration types and face-selective regions. On average, a substantial proportion of vertices in each face-selective area were active during face perception and encoding (Figure 2g). Large proportions of activated vertices directly confirmed the overlap we noticed based on visual inspection. Second, to compare the activation magnitude between face perception and encoding, we computed the average t-statistic values for each face-selective region. Figure 2f depicts the t-statistic values for the three face-selective areas between perception and encoding. Group linear mixed-effect analysis of activation magnitude, with task (perception/encoding), face-selective region (mFus, pFus, IOG) and hemisphere (left and right) as fixed effects and subject as random effects, revealed significant main effects of task and ROI, but no significant difference between left and right hemisphere ROIs (Satterthwaite approximations used for significance of model coefficients). Specifically, using pairwise Tukey’s range test, with p-values adjusted for comparing a family of 3 estimates, activation magnitude was greater for encoding than perception (encoding - perception: *t*(154) = 5.314, p <.0001); and IOG and pFus activation were greater than mFus (IOG – mFus: *t*(156) = 4.323, p = .0001; pFus – mFus: *t*(155) = 2.370, p = 0.496). This indicates that while there was reduced overlap between perception and memory encoding, the activity magnitude was greater during memory encoding than perception. Together, these data confirmed that identified face selective cortex, within each subject, served as a robust means for identifying the functional substrate engaged during face stimulus memory encoding.

In summary, we successfully mapped the unique pattern of face-selective regions in each participant’s VTC via a separate controlled visual object localizer task. Our data suggested systematic differences between individuals in the number and configuration of face-selective regions, consistent with recent population analyses on VTC selectivity mapping (Chen et al., 2023). Despite these variations, we found that face encoding activity during the study phase of a memory paradigm engaged similar brain regions that were identified as face-selective at an individual level. While this direct comparison between perception and memory encoding further supported the notion that cortical regions engaged in perception are also active during visual encoding, specifically, visual category stimuli engage their visual-selective regions for information processing (Bainbridge et al., 2021; Rugg & Thompson-Schill, 2013; Silson et al., 2019; Srokova et al., 2022; Steel et al., 2021), this large functional overlap sets up the stage for our next questions regarding cortical regions involved in memory retrieval and their functional overlap with encoding and perception.

### Memory reactivation in face-selective regions

To examine evidence of cortical reinstatement, we examined whether regions in VTC that were engaged during memory retrieval overlapped with the brain regions involved in perception and memory encoding. Here, we examined the responses during memory retrieval using the same approach outlined above for memory encoding analysis. During the memory retrieval phase, participants were presented with cue words alone without associated face images. Participants were asked to vividly retrieve faces that were previously paired with the old cue words during the encoding phase (Figure 1b). As external visual inputs of face stimuli were absent during retrieval, the neural responses in the identified face-selective regions would likely be driven by internal processes such as memory reinstatement rather than external visual processing. Using GLM contrasts of successful retrieval of old (hits) > new (correct rejections), we identified memory retrieval activation maps that revealed the brain areas involved in successful retrieval of cue-associated faces. Figure 3a illustrates the memory retrieval activity maps for the same three example configuration types as in Figure 2, with putative face-selective regions outlined in black on the inflated cortical surface. These activation maps were generated using a common threshold across participants (t > 2, vertex level). Given the absence of external face stimuli input during retrieval, we expected reduced VTC activity compared to perceptual encoding. However, we still anticipated that face-selective regions would be active to support memory retrieval of faces. Visually, VTC engagement during memory retrieval appeared less extensive compared to encoding (Figure 2).

**Figure 3.**
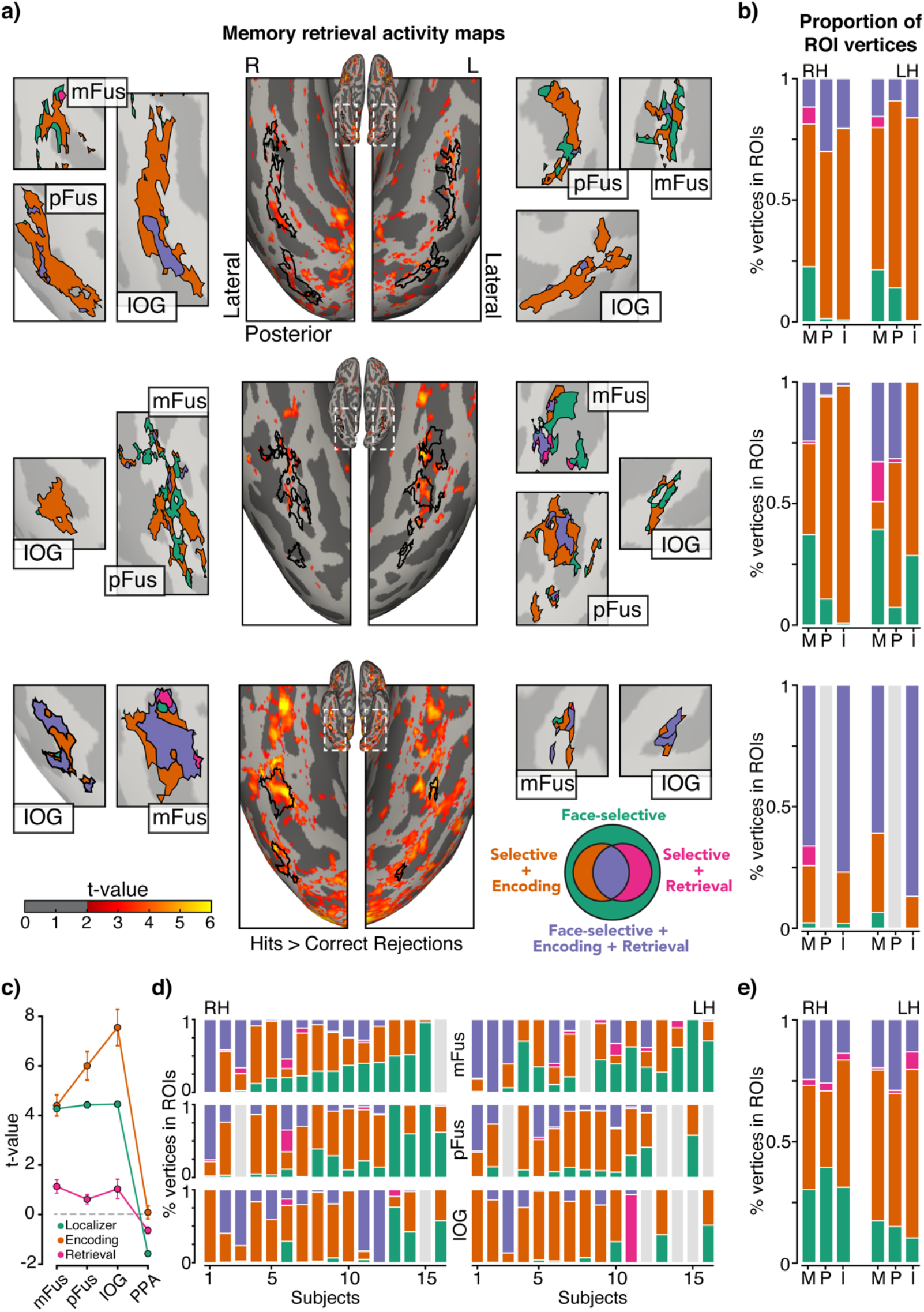
Memory retrieval activity within ventral temporal cortex face-selective regions. **a)** Example participants showing Memory retrieval activities in the ventral temporal cortex (center, GLM contrast map: hits > correct rejections) for the separate (top), continuous (middle), and single (bottom) type example participants in (Figure 2a). Black outlines denote the boundaries of face-selective regions identified in Figure 2a. Color maps illustrate the t-statistic of activation of the contrast. Zoomed-in maps of each face-selective region illustrate the spatial distribution of vertices within the region that were active for i) face-selective localizer only (green), ii) localizer and memory encoding (orange), iii) localizer and memory retrieval (magenta) and iv) localizer, memory encoding, and retrieval (purple) in the left and right hemisphere. **b)** Proportions of vertices within each face-selective region (mFus(M), pFus(P) and IOG(I)) that were active for the same example participants during the same contrasts in (a). **c)** Mean t-value of GLM contrasts (Localizer: Face > other visual categories; Memory Encoding: Stimulus presentation > inter-trial fixation and Memory retrieval: hits > correct rejections) are shown across three face-selective regions (mFus, pFus and IOG) and a control region (parahippocampal place area, PPA) averaged between left and right hemispheres. **d)** Proportions of active vertices are shown for all participants across three face-selective regions in both left and right hemispheres. **e)** Group averaged proportions of active vertices in each face-selective region.

To quantify the spatial overlap of responses, we followed a similar approach to our memory encoding analysis. First, using face-selective regions as anatomical masks, we identified vertices as active during memory retrieval if GLM contrast t-statistic was greater than 2. Figure 3b shows the proportions of active vertices during different scenarios of overlap: perception only, perception and encoding, perception and retrieval, or perception, encoding, and retrieval, for the three example configuration types and face-selective regions. Additionally, the spatial distribution of active vertices within regions for different overlap scenarios is shown in Figure 3a. On average, approximately 25% of vertices were active during face perception, memory encoding, and memory retrieval across all face-selective areas (Figure 3e). This result indicates that portions of these face-selective regions are reactivated, with retrieval activity conforming to individual perception/encoding neural substrates, supporting the notion of cortical reinstatement. Moreover, only a small number of vertices were active during retrieval but not during encoding (< 5% across participants), indicating a high degree of encoding-retrieval overlap. It is worth noting that the perception-encoding-retrieval overlap varied considerably across regions and participants (as shown in Figure 3d). For example, while the separate configuration type participant (Figure 3b top) had around 20% of vertices overlap during face perception, memory encoding, and memory retrieval, the single configuration type participant (Figure 3b bottom) had more than 50% of vertices overlap. Given these differences, we examined the relationship between the extent of individual activity overlap across perception-encoding-retrieval conditions and memory performance (*d’)*. Consistent with prior observations, we observed greater response overlap to be significantly correlated with improved memory performance (Spearman 𝜌 = 0.76, p <0.001; n =16; 𝜌 95% c.i.: 0.52:0.94).

Next, we compared the activation magnitude during face perception, memory encoding, and retrieval in face-selective regions to a control brain region, which is a common approach to assess the degree of category selectivity (i.e., face vs. place). We expected that successful retrieval of face stimuli would elicit a response in face-selective regions but not in place-selective regions. Alternatively, if the responses were similar between face and place-selective regions, it may suggest that memory retrieval reactivation of cortical regions was not content-specific, likely a more general top-down modulation of sensory areas (Kastner et al., 1999; Righart et al., 2010). Therefore, we localized the parahippocampal place area (PPA) in each participant using a similar approach as used for identifying face-selective regions. Briefly, we used a GLM contrast to identify regions where responses to corridor/house stimuli were greater than eight other visual categories shown (face, bodies, cars, instruments, limbs, letters, numbers, and phase-scrambled noise). Place-selective clusters were identified based on a common t-statistic threshold across participants (t>3, vertex level). From the place-selective clusters, PPA was identified based on their anatomical location (Epstein et al., 1999) in every participant (n=16). The average t-statistic values for each face-selective region and PPA during perception, memory encoding, and memory retrieval were computed and depicted in Figure 3c. Extending our group linear mixed-effect analysis of activation magnitude, with task (perception/encoding/retrieval), regions of interest (mFus, pFus, IOG, PPA) and hemisphere (left and right) as fixed effects and subject as random effects, revealed significant main effects of task and ROI, but no significant effect between left and right hemisphere ROIs (Satterthwaite approximations used for significance of model coefficients). Specifically, using pairwise Tukey’s range test, with p-values adjusted for comparing a family of 3 estimates, activation magnitude during retrieval was substantially reduced (Perception – Retrieval: *t*(335) = 8.910 p<.0001; Encoding – Retrieval: *t*(335) = 15.226, p < .0001).

In addition, activation magnitude was greater in face-selective regions compared to PPA (mFus – PPA: *t*(335)=13.820, p<.0001; pFus – PPA: *t*(337)= 14.891, p <.0001; IOG – PPA: *t*(337) =17.163, p<.0001). As noted above, this reinstatement activity is predicted to closely recapitulate patterns observed during encoding, although with a reduced magnitude when compared to sensory-driven responses. When compared to PPA, the face-selective regions were more activated during face perception, and memory encoding, supporting their role in processing face stimuli.

In summary, using individually mapped face-selective regions in each participant’s VTC, we found that face retrieval activity engaged an overlapping subset of cortical vertices. The engagement of these face-selective regions was reduced during retrieval compared to face perception and encoding but was still more activated when compared to a place-selective control region. Overall, our study adds to the growing body of evidence supporting the cortical reinstatement theory of memory retrieval and indicates that memory reinstatement is an information-specific process, where the precise brain regions involved in processing specific stimuli during encoding are reactivated during successful memory retrieval.

### Memory transformation in face-selective regions

Our results showed that cortical activation overlap between face perception and memory encoding was substantial, with approximately 75% of face-selective vertices showing activity in both processes. However, the overlap between face perception/encoding and memory retrieval was more modest, with around 25% of face-selective vertices being active during retrieval. This may indicate that only a subset of face-selective regions was activated because memory retrieval involves selective reactivation of specific neural networks associated with the initial sensory-perceptual experience (Bone et al., 2020). One possible alternative explanation for this modest retrieval activity level and overlap is the “spatial transformation” hypothesis, which posits that the cortical spatial location of neural activity during memory retrieval systemically differs from that of perception and encoding. This idea is supported by evidence from studies of place/scene-processing networks that have observed a systematic “anterior shift” in cortical regions involved in scene retrieval compared to those involved in scene perception or memory encoding (Bainbridge et al., 2021; Baldassano et al., 2016; Fairhall & Ishai, 2007; Rugg & Thompson-Schill, 2013; Silson et al., 2016; Steel et al., 2021). To test whether a similar anterior shift occurs in individual face-selective regions during memory retrieval, we examined the location of neural responses during perception and compared them to the activity locations during memory encoding and retrieval.

To quantify any activity shift within VTC, we employed the weighted center of mass approach outlined in Steel et al.,(2021) combined with individually identified face-selective regions, considering both the activation level and cortical geometry of regions identified. We first extracted activity maps using the identified face-selective regions as masks during face perception, memory encoding, and retrieval. Then we calculated the center of mass weighted by their respective activation (Figure 4a shows an example participant). The zoomed-in maps in Figure 4a display the activity-weighted center of mass for each face-selective region. The centers of mass for face perception, memory encoding, and retrieval appeared to be very close in their anatomical locations. Therefore, to better quantify and visualize the possible location shift, we represented these centers of mass as coordinates on a polar plot, with the x-axis representing the medial-lateral direction and the y-axis representing the anterior-posterior direction. In this polar plot, unity (0,0) was defined by the location of the center of mass weighted by face-perception activity for all three face-selective regions (as shown in Figure 4b). For the example participant, the centers of mass weighted by encoding activity (orange color) and retrieval activity (magenta color) appeared to be shifted away from the center of mass weighted by face perception activity. However, the observed changes were relatively small, all within 2 mm in distance. As a point of reference, the size of anterior shifts during scene retrieval has been reported to range from 10-30 mm, relative to encoding substrates (Silson et al., 2019; Srokova et al., 2022; Steel et al., 2021). Most importantly, there was no clear systematic shift in any direction.

**Figure 4.**
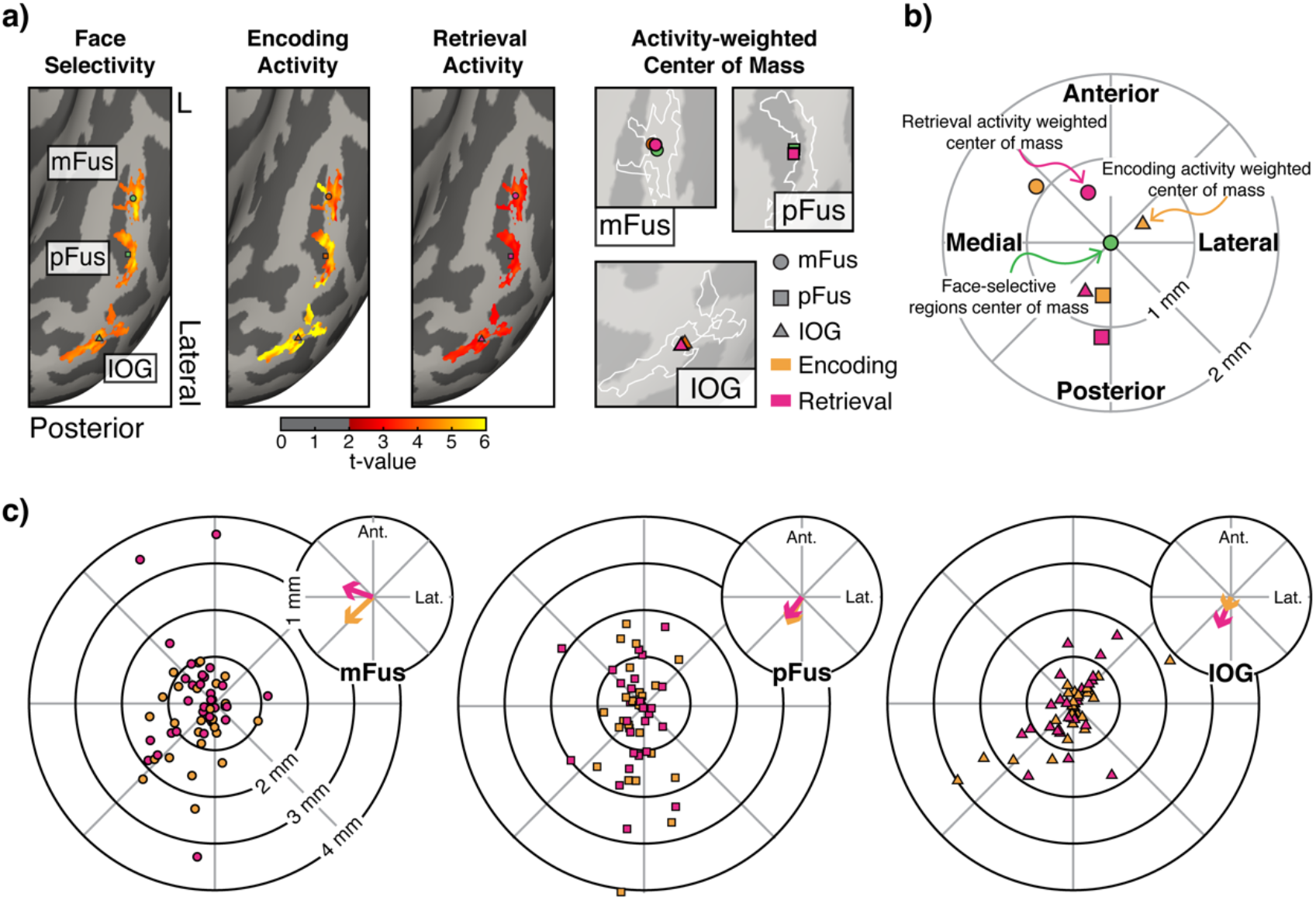
Activity-weighted center locations of face selectivity, memory encoding, and retrieval activity within ventral temporal cortex face-selective areas. **a)** Example participant showing activity in the VTC within the boundaries of face-selective regions identified in Figure 2a during object localizer (GLM contrast map: face > all other visual categories), memory encoding (GLM contrast map: stimulus presentation > fixation), and memory retrieval (GLM contrast map: hits > correct rejections). Color maps illustrate the t-statistic of activation of the contrast. Zoomed-in maps of each face-selective region illustrate the activity-weighted center of mass. White outlines denote the boundaries of face-selective regions. **b)** Example participant summary polar plot showing topography of memory encoding and retrieval activity relative to face selectivity. The activity-weighted center of mass for face selectivity (green) in three face-selective regions is at unity (0,0). The centers of mass for memory encoding (orange) and retrieval (magenta) responses are plotted in relation to face selectivity for mFus (circle), pFus (square) and IOG (triangle). X-axis represents the medial-lateral direction and the y-axis represents the anterior-posterior direction. **c)** Topography of memory encoding and retrieval activity relative to face-selectivity are shown for all participants across three face-selective regions. The centers of mass for memory encoding and retrieval activity are plotted in relation to the center of mass for face selectivity (unity, same as in b). Zoomed-in maps of each plot illustrate the averaged vector distance and direction of shift in the center of mass for memory encoding (orange) and memory retrieval (magenta) activity.

A similar pattern emerged when we examined the activity-weighted centers of mass for all participants. Figure 4c provides summary polar plots for the center of mass positions in the three face-selective regions (similar to the example participant in Figure 4b). Overall, most shifts during memory retrieval were concentrated within a 2 mm distance from the center of mass for face perception and did not display a particular directional bias. More precisely, when we calculated the average shift in distance and direction (see Figure 4c zoomed-in maps), we found that the average distance shift for encoding and retrieval activity from face perception was less than 0.5 mm. Again, there was no consistent preferred direction of shifts in activity.

In summary, our results demonstrate that the anatomical locations of face-selective regions during memory retrieval did not significantly differ from those observed during face perception and memory encoding. When comparing the brain activity patterns during memory encoding and retrieval, we consistently observed a reduction in spatial overlap and magnitude of responses. However, the specific location within the brain where this reduced activity occurred remained similar to the location observed during face perception. This lack of significant spatial shift between perception and memory retrieval suggests that spatial transformation, involving large-scale shifts in cortical activations, is not a prominent feature during the retrieval of face stimuli. Instead, the brain regions that are activated during perception of faces are also involved in the retrieval of face memories, albeit with reduced activity levels.

### Electrophysiological recordings from face-selective regions during perception, memory encoding, and retrieval

In order to better understand the neurophysiological basis of our fMRI findings and to examine the temporal dynamics of sensory reinstatement we conducted a human intracranial electrocorticography (ECoG) experiment. Specifically, we had the opportunity to perform ECoG recordings from a rare patient with refractory epilepsy who underwent invasive monitoring with electrodes placed directly on VTC, uniquely covering a large portion of the left fusiform gyrus (See Figure 5c). To align ECoG data with our fMRI findings presented above, the patient performed identical visual object localizer and word-face memory tasks (Figure 5a,b).

**Figure 5.**
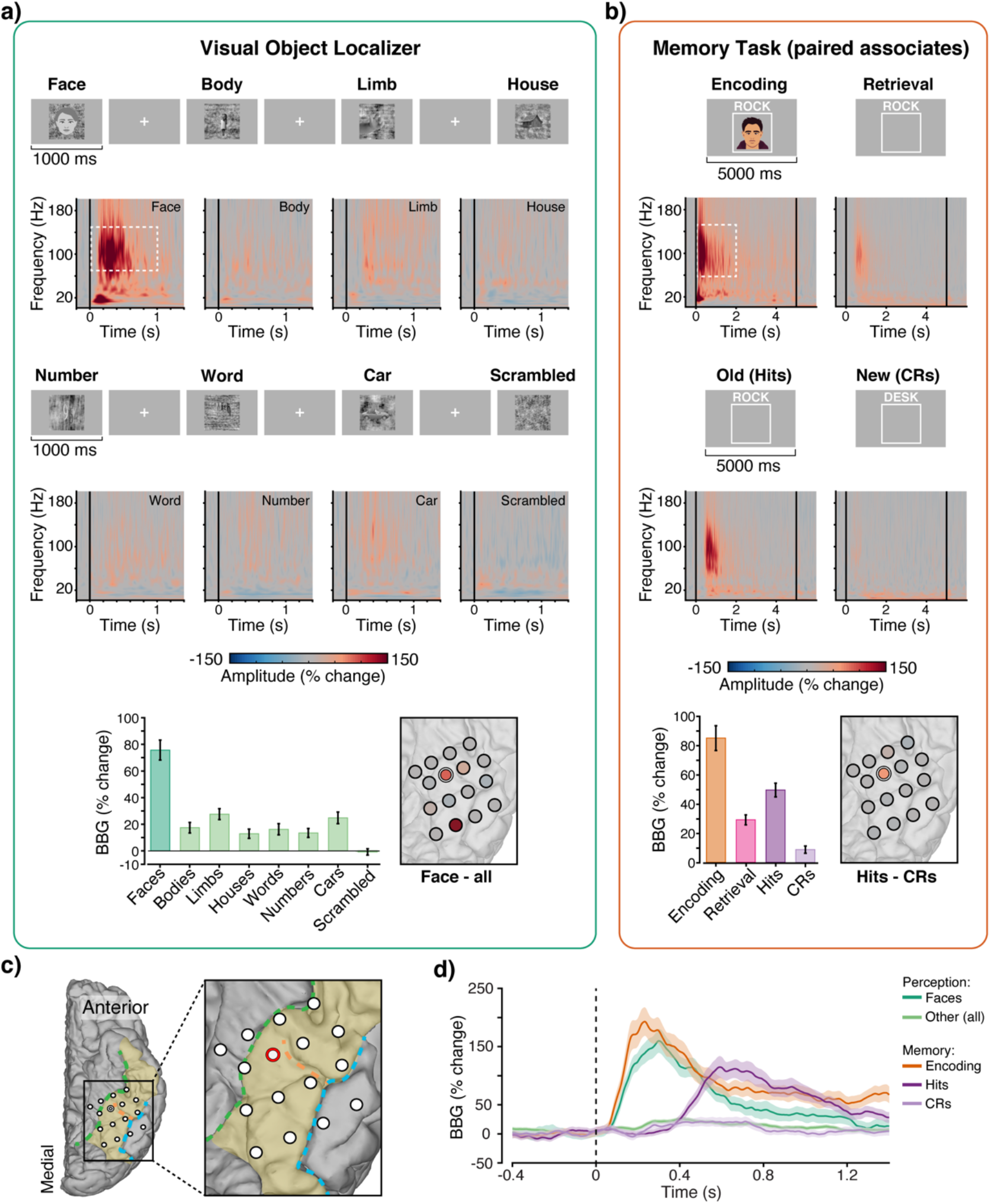
ECoG recordings from the ventral temporal cortex during face perception, memory encoding, and retrieval. **a)** Visual object localizer task for identifying face-selective areas. Greyscale images from eight visual categories were presented for 1,000 ms with a random ISI for 1-1.5 seconds. Patient had to provide a button press to indicate any 1-back stimulus repetition (Same experimental procedure and stimuli as fMRI, Figure 1). Example stimuli from each of the eight visual categories as shown. Average spectrograms from one example electrode (indicated in b) are shown for each visual category; color maps reflect percentage changes in amplitude relative to the pre-stimulus period (the black line indicates stimulus presentation onset). The mean amplitude response of broadband gamma activity (BBG, 70 – 150 Hz) during stimulus presentation (0 – 1000 ms) are shown for all eight visual categories (bottom left). The white dashed-lined box in the spectrogram denotes the frequency range and time window used to extract broadband gamma (BBG; 70-150 Hz) activity for response calculations. Error bars reflect SEM. These plots show that this electrode location displays strong face selectivity. The color of each electrode denotes the mean BBG response contrast, defined by the BBG response to face stimuli minus the response to all other visual categories combined (Face – all). Warmer colors indicate stronger face selectivity. **b)** Memory experimental procedure and mean spectrogram from one example electrode (same as a). During the encoding phase, word-image pairs were presented for 5,000 ms with a self-paced inter-stimulus interval (ISI). During the retrieval phase, cue words (old/studied and new/unstudied words) were presented for 5,000 ms followed by a memory strength judgment. Patient was required to encode word-face associations and to retrieve the associated face image from previously studied word cues. Average spectrograms are shown for memory encoding and retrieval. Retrieval was further separated based on cue type and behavior: hits and correct rejections; color maps reflect percentage changes in amplitude relative to the pre-stimulus period (the black line indicates stimulus presentation onset). The mean amplitude response of BBG during stimulus presentation (0 – 2000 ms) are shown for encoding, retrieval, hits and correct rejections (bottom left). The white dashed-lined box in the spectrogram denotes the frequency range and time window used in BBG activity response calculation. Error bars reflect SEM. These plots show that this electrode location was active during encoding and retrieval, specifically during the successful retrieval of old word-face pairs (hits). The color of each electrode denotes the mean BBG response contrast defined by the BBG response to hits minus the response to correct rejections (Hits – Correct Rejects). Warmer colors indicate successful memory retrieval. **c)** Anatomical locations of ventral temporal electrodes in one patient with the example electrode outlined in red. Major ventral temporal anatomical landmarks colored: fusiform gyrus (yellow), collateral sulcus (green), mid-fusiform sulcus (orange), and occipitotemporal sulcus(blue). **d)** Average mean amplitude time course for BBG ranges (shading reflects SEM) for perception (face and other visual categories) and memory (encoding, hits, and correct rejections).

First, using time-frequency analysis, we determined visual selectivity by examining the spectral response to each presented visual category (Figure 5a, top) and then calculated the average broadband gamma response (BBG, 70 - 150 Hz) (Bartoli et al., 2019). Face stimuli elicited higher BBG responses than all other categories, indicating face selectivity (Figure 5a, bottom left). BBG analysis across VTC electrodes revealed two face selective sites, likely recording from mFus (example electrode, outlined in white) and pFus. As electrodes between these face-selective sites lacked face-selective responses, this patient was classed as having a “separate” type configuration. Next, we examined the extent to which these face-selective regions were electrophysiologically engaged during memory encoding and retrieval. As with our fMRI study, the patient performed a memory task involving word-face pairs, and later, retrieved associated faces when given word cues (Figure 5b). During memory encoding, we expected that face images would elicit strong BBG responses within identified face-selective sites. Indeed, as expected, face-selective electrodes showed clear responses during encoding (Figure 5b top left), confirming activity overlap between perception and memory encoding, consistent with our fMRI finding.

During memory retrieval, the patient was presented with cue words without associated face images. Notably, increased BBG activity was observed in response to cue words, although with a smaller response magnitude than at encoding. Strikingly, when retrieval responses were separated by hits (old) and correct rejections (new), BBG responses were substantially larger for hits (Figure 5b middle). To examine the spatial transformation hypothesis, we investigated whether the location of neural activity during memory retrieval shifted compared to perception and encoding. Notably, aside from the example electrode site (outlined in white), no other electrodes exhibited a robust memory retrieval effect (Figure 5b, bottom right). The focal nature of this memory retrieval effect aligns with memory reinstatement theory, consistent with our fMRI findings.

Finally, it is important to highlight the difference in the onset of BBG activity between memory encoding and retrieval, not typically resolvable with fMRI. During encoding, BBG activity showed a rapid onset within 100 ms after stimulus presentation. This suggests that increased BBG activity was associated with externally driven visual processing that quickly responds to the presented face stimuli. In contrast, during memory retrieval, BBG activity displayed delayed onset at around 400 ms after the presentation of cue words (Figure 5d). This difference in the onset implies that while memory retrieval neural activity occurs in the same cortical region as encoding, it undergoes a temporal shift – delayed during retrieval compared to encoding’s rapid response.

## Discussion

In this study, we aimed to investigate the organization, and potential transformation, of activity patterns within the human VTC during the perception, encoding, and retrieval of face stimuli. To do so, we conducted a multi-session fMRI study to identify individual-specific responses across these conditions. Through this individualized approach, we tested how closely memory-driven reinstatement activity recapitulates individual specific perceptual/encoding substrates and quantified any consistent topographic transformations in activity patterns across these conditions. Our results showed that individuals displayed common types of variability in the size and configuration of face-selective regions, consistent with recent large-scale findings. We also observed a significant overlap between activation patterns during face perception and memory encoding, specific to each individual. During memory retrieval, there was a reduction in activation levels of face-selective regions compared to face perception and encoding. Importantly, we did not find evidence of a consistent topographical shift in the anatomical location of face-selective regions during memory retrieval compared to perception and encoding, indicating the absence of spatial transformation. Furthermore, our direct electrophysiological recordings from the human VTC, under the same experimental conditions, demonstrated similar spatial effects, but revealed a clear change in the timing of neural activity onset between encoding and retrieval. Our study provides evidence supporting the view that retrieval-driven cortical reinstatement closely corresponds to individual-specific encoding substrates. However, in contrast to recent findings in scene perception and memory, we did not observe consistent spatial transformation in activity patterns between perception and memory for face stimuli. This suggests that the anteriorization of retrieval activity and its underlying neural mechanisms may not apply universally to other visual categories. Our results highlight the importance of assessing functional organization at an individual level when studying memory reinstatement and examining other forms of memory transformations from encoding to retrieval.

Using controlled face stimuli and a separate visual object localizer experiment, we identified three distinct face-selective regions (mFus, pFus, and IOG) for each participant. These regions were categorized into three configuration types: “separate,” “continuous,” and “single”. Despite having a more limited sample size, our regional delineation was consistent with previous studies on the functional organization of VTC (Julian et al., 2012; McGugin et al., 2016; Pinsk et al., 2009; Weiner & Grill-Spector, 2010, 2012, 2013). Furthermore, the proportions of configuration types we observed closely matched those reported in a large-scale study involving 1,000 participants that specifically focused on face-selective regions (Chen et al., 2023). The individual differences in face-selective configuration types were further supported by our observation of a substantial overlap between the cortical areas engaged during face perception and memory encoding. Historically, individual variability has not been given sufficient consideration in prior studies that explore the retrieval responses for face stimuli. For example, some studies have focused on activities within a singular FFA region, while others included activity from a broader area, such as the entire fusiform gyrus or VTC (Cichy et al., 2011; Haxby et al., 2001; Kuskowski & Pardo, 1999; O’Toole et al., 2005; Prince et al., 2009). Functional organization of VTC is complex, with multiple visual-category selective regions, including regions responsible for processing faces, limbs, and bodies within the fusiform gyrus alone (Weiner & Grill-Spector, 2010). This complex organization is important to consider when examining group-level findings, together with spatial smoothing, which may blur or combine distinct face-selective regions, or even incorporate non-face-selective regions (Weiner & Grill-Spector, 2012). Furthermore, it’s worth noting that in addition to the impacts of certain analysis approaches, distortion in the configuration of identified face-selective maps is also sensitive to the criteria or contrast employed to capture face selectivity. Previous studies have frequently defined face-selective regions based on contrasts of face vs. place condition stimuli during encoding, or used a priori anatomical masks from brain atlases, which may overestimate or misrepresent VTC regions specialized for face stimuli (Schwarz et al., 2019). In contrast, our focus on identifying individual-specific response patterns with stringent selectivity criteria (i.e., face vs. multiple other visual categories) allowed us to provide a more precise examination of reinstatement properties by considering the individual-specific encoding substrates of each participant.

Our investigation of the overlap between face perception/encoding and memory retrieval demonstrated that a subset of face-selective vertices was active during retrieval, as predicted. This finding suggests that memory retrieval involves the selective reactivation of specific neural networks associated with the initial sensory-perceptual experience, adding support to cortical reinstatement theory. Earlier fMRI studies have also observed similar patterns, with sensory cortices engaged during encoding showing partial reactivation during retrieval (Wheeler et al., 2000, see, Danker & Anderson, 2010, for a review). However, careful inspection of the literature shows conflicting results with the tenants of retrieval-driven reinstatement. For example, several studies report on category reinstatement within regions typically not associated as being category selective, or do not closely overlap with identified encoding substrates (McDermott et al., 1999; Moscovitch et al., 1995; Vaidya et al., 2002). More broadly, some approaches cast a broad anatomical region of interest, limiting the ability to assess the spatial precision of reinstatement (Rissman & Wagner, 2012; Ritchey et al., 2013; Rugg et al., 2008). Our approach allowed us to examine activity within each ROI at the vertex level. We found that approximately 25% of face-selective vertices were reactivated during retrieval, indicating partial reactivation happening within each ROI. The notion of partial reactivation is in line with the concept of hippocampal “pattern completion” processes, where neural representations do not need to be perfectly reactivated for successful retrieval (Danker et al., 2016; Horner et al., 2015; Rugg et al., 2008). This difference in neural representation may be necessary to help distinguish between a current perceptual experience and the active recall of a prior perceptual experience. Lastly, an interesting observation is that the extent of overlap between perception, encoding, and retrieval processes displayed a positive correlation with memory performance across participants. This correlation aligns closely with prior research that has examined encoding-retrieval similarity across diverse brain regions and found that greater similarity in neural activity patterns between the encoding and retrieval phases corresponds with more effective memory outcomes (Favila et al., 2020).

Memory retrieval activity is not a perfect recapitulation of perception and encoding. A growing literature has focused on understanding the transformations of neural representation between perception and memory retrieval (Favila et al., 2020). Topographic spatial transformation has been chiefly supported by research on place/scene-processing networks, which has shown an anterior shift in cortical regions during scene retrieval compared to perception and encoding (Baldassano et al., 2016; Fairhall & Ishai, 2007; Silson et al., 2016, 2019; Srokova et al., 2022; Steel et al., 2021). However, in our investigation, when carefully considering individual-specific face-selective regions, we did not observe a similar anterior shift during memory retrieval. There could be several reasons for this discrepancy. One possibility, as mentioned above, may be the result of group-level analysis. However, Steel et al. (2021) conducted individual-level analyses and still observed an anterior shift in PPA activity during memory retrieval compared to perception. Interestingly, in the same study, Steel et al. (2021) did not find an anterior shift for face stimuli in FFA. It is possible that the delineation of a single FFA might contribute to the absence of the shift. In our study, we carefully assessed the locations of perception and retrieval activity with face-selective regions identified for each individual but still did not find a consistent spatial transformation. Therefore, the anterior shift may be specific to the scene-processing network. Steel et al. (2021) suggested that the PPA’s neural representations exhibit an anterior-posterior gradient, shifting from concrete/perceptual during perception and encoding to more abstract/conceptual during memory retrieval. This gradient may arise from functional connectivity differences, as anterior PPA is more connected to the default mode network (DMN), supporting memory retrieval, while posterior PPA is more connected to occipital visual regions associated with perception (Baldassano et al., 2013, 2016; Peer et al., 2019; Silson et al., 2019). Although these studies provide some theoretical basis for the anteriorization of scene-related memory, it remains unclear why this gradient does not apply to other visual categories, and how it can accommodate the complex interdigitated organization of category selectivity within VTC.

To complement our fMRI results, we conducted the same experiments on a patient with VTC ECoG recording, providing high temporal resolution and anatomical precision. The ECoG findings supported our fMRI results, showing activity in the same face-selective regions during encoding as during visual perception. During face retrieval, specific face-selective regions were reactivated, indicating focal memory reinstatement. Importantly, we did not observe a clear spatial shift in memory retrieval activity, reinforcing the idea of localized cortical reinstatement rather than a spatial transformation. While many studies have focused on spatial memory transformations, our ECoG results revealed a temporal shift in neural activity during memory retrieval compared to encoding. This change in temporal dynamics likely reflects the inherent distinction between perceptual experiences and memories. Specifically, the delayed response during retrieval aligns with the concept that the hippocampus engages in pattern completion processes, triggering targeted sensory cortices to reinstate neural representations for memory retrieval. Typically, this sensory reinstatement occurs around 500 ms after the cue onset, consistent with our findings (see Staresina & Wimber, 2019, for a review). Therefore, our results emphasize the importance of considering transformation in the temporal domain when studying memory retrieval processes. Indeed, temporal transformation has been observed in rodents, where the firing of place cells occurs faster during sleep compared to maze exploration, indicating “temporal compression” (Bosch et al., 2014; Ólafsdóttir et al., 2018). In humans, electrophysiological recordings during rest or sleep also suggest a similar compression of memory replay (Liu et al., 2019; Michelmann et al., 2018). Understanding these temporal dynamic principles of memory retrieval can provide valuable insights into the neural order of operations underlying remembering.

Our individualized approach allowed us to carefully investigate the involvement of face-selective regions in VTC concerning memory reinstatement and transformation of face stimuli. However, there are some limitations to our study. First, the face stimuli used in the visual object localizer and memory task had differences in perceptual features (gray-scale images vs. colored images) and familiarity (unfamiliar faces vs. famous faces). These variations in face stimuli used for perception and memory tasks may have introduced variance in neural representations, which could potentially explain why the activation overlap between these two conditions was not an exact match and why the magnitude of activation differed. Additionally, our fMRI sample size was relatively small, and the electrophysiology recording was limited to a single-patient case study. While a small sample size may limit the generalizability of findings to the larger population, it is worth noting that we observed similar proportions of face-selective region configuration types as those reported in the large-scale study of face selectivity organization (Chen et al., 2023). Moreover, our analyses were constrained to individually identified face-selective regions, which could potentially overlook activations or interactions in other brain areas. However, when we compared face-selective regions to the place/scene-selective PPA as a control region, the face-selective regions exhibited significantly higher activity during retrieval. This observation suggests that the retrieval process is category-specific and focal to face-selective regions. While our approach was relatively conservative in ROI selection, it ensured that the retrieval activity was specific to face-related memory. Further research is necessary to explore memory reinstatement and transformation in different brain regions and other visual categories. This would allow for a more comprehensive understanding of the underlying neural processes involved in memory retrieval.

In summary, our study used an individualized approach, identifying specific responses for each participant to investigate memory-driven reinstatement and consistent topographic transformations in activity patterns. Our findings revealed that retrieval-driven cortical reinstatement closely aligns with individual-specific encoding substrates. Importantly, we did not observe consistent spatial shifts in face-selective regions between perception and memory, inconsistent with findings from studies of scene-selective regions. Electrophysiological recordings from the human VTC further supported our findings, but highlighted important changes in the timing of neural activity onset between perception and memory. Our results underscore the importance of assessing functional organization at an individual level, as well as considering the temporal domain, in future efforts to elucidate the principles of perception-memory transformations in the human brain.

## Methods

### Participants

Sixteen healthy participants (7 females, 2 self-reported left-handers, mean age 27.9 yrs., ranging from 18 – 39 yrs.) with normal or corrected to normal vision completed the experiment. Informed consent was obtained prior to participation. The experimental protocol was approved by the Committee for the Protection of Human Subjects of the Baylor College of Medicine, Houston, TX (IRB protocol number H-38398). All participants were monetarily compensated for their time.

### fMRI Experimental Design

To ensure a more robust neural response and limit the fatigue of participants, this study comprises two experimental sessions. Session 1 was a 60-minute scanning session of two high-resolution anatomical structural scans, three runs of the visual object localizer task for robust identification of visual category selectivity in the ventral temporal cortex. Session 2 took place on average 39 hours (1.6 days, ranging from 22 – 90 hrs.) after Session 1. It was a 90-minute scanning session of one high-resolution anatomical structural scan and three runs of the paired associates memory task. Participants were scanned at Baylor College of Medicine’s Core for Advanced MRI (CAMRI). Stimuli were presented on an MR compatible 32” LCD Display screen (BOLDscreen, Cambridge Research Systems [CRS], Rochester, UK) placed behind the bore of the scanner. Behavioral responses were collected using a fiber-optic button response pad (Current Designs, Haverford, PA, USA). Stimuli were presented and synchronized with the MR data acquisition using MATLAB (The Mathworks, Inc., Natick, MA, USA) and the Psychophysics Toolbox (Brainard, 1997; Pelli, 1997).

#### Session 1. Visual object localizer

During the visual object localizer task, participants were presented with grayscale images from 10 visual categories: faces, bodies, cars, corridors, instruments, limbs, houses, letters, numbers, and phase-scrambled noise. Visual stimuli come from a publicly available corpus that has been successfully used as a visual category localizer in fMRI studies (Stigliani et al., 2015). The stimuli have been statistically matched for contrast, luminance similarity, and spatial frequency. Following prior work, stimuli were presented in a mini-block design. During each mini-block, participants viewed eight images from the same visual category at a rate of 500 ms per image, for a total of 4,000 ms (Figure 1a). A total of 60 mini-blocks (6 mini-blocks per visual category) were presented in random order. Across the entire experiment, 20% of the mini blocks (12 mini-blocks) contained a target trial (repeat images), where participants were asked to respond via button press when an identical stimulus was repeated back-to-back (1-back task, Figure 1a). Participants performed 3 runs of the visual object localizer task (about 6 min per run) to allow robust identification of visual category selectivity in the ventral temporal cortex.

#### Session 2. Paired associates memory task

Visual object localizer and memory tasks were scanned in separate sessions to help reduce participant fatigue from long scanning time without compromising data quality. Localizer and memory tasks were conducted on average 1.6 days apart. There were three phases to the memory task: encoding, delay, and retrieval. During the encoding phase, participants were presented with 30 word-picture pairs. For each trial, single words (e.g., ‘ROCK’, ‘GLASS’) were displayed above color photographs of well-known people (e.g., ‘Barack Obama’ or ‘Taylor Swift’) for 5 seconds per pair with a jittered inter-stimulus interval between 4-6 seconds (Figure 1b). Participants were instructed to associate each arbitrary word-face pair. After encoding, during the delay phase, participants were presented with 14 trials of a single-digit number (e.g., ‘6’ or ‘5’) and were asked to judge whether the number was odd or even via button press. Numbers were presented for 1500 ms with 500 ms inter-stimulus interval. During the retrieval phase, participants were presented with 60 cue words (30 old/studied and 30 new/unstudied, not associated with a face image). Single cue words were presented for 5 seconds, during which participants were instructed to retrieve and “bring to mind” the associated face picture as vividly as possible. Once the cue was removed, participants were asked to rate their memory via a button press on a 4-point scale: 1) “No memory”; 2) “Yes, memory for word only”; 3) “Yes, memory for word and picture weak”; 4) “Yes, memory for word and picture strong”. The question/response screen was presented for 5 seconds with a jittered inter-stimulus interval between 4-6 seconds (Figure 1b). Participants performed 3 runs of the memory task (about 21 min per run) to allow robust identification of memory encoding and retrieval activity in the ventral temporal cortex.

Word stimuli are selected with a limited letter range (4-8), number of syllables (1-3), concreteness rating (600-700), and imaginability rating (600-700) from the Medical Research Council Psycholinguistic Database (http://www.psy.uwa.edu.au/MRCDataBase/uwa_mrc.htm). Face stimuli were color images of famous individuals with an equal number of women and men from Google Image Search. Images were rescaled to 450 X 450 pixels, with a resolution of 97.987 pixels/inch.

### fMRI methods

#### Imaging acquisition

Imaging data were acquired using a 3T Siemens Trio MRI scanner (Siemens AG, Erlangen, Germany) equipped with a 32-channel head coil. A total of three high-resolution T1-weighted anatomical scans (TR = 2600 ms, matrix = 256 x 256 mm^2^, 176 slices, voxel size = 1.0 X 1.0 X 1.0 mm^3^; FOV = 256 mm, TE = 3.02 ms; flip angle = 8°) were collected. Of these, two T1 scans were collected at the beginning of Session 1 before the visual object localizer, and one scan was collected at the beginning of Session 2 before the memory task. The functional MRI scans covered the entire brain using a continuous multi-slice echo planar sequence. Scanning parameters were identical across functional scans (visual object localizer, resting state, and memory task): TR = 2000 ms, TE = 30 ms, flip angle = 72°, in-plane resolution of 2 x 2 mm, 69 2 mm axial slices, multiband acceleration factor = 3, voxel size = 2.0 X 2.0 X 2.0 mm^3^. A total of 153 volumes acquired for each run of visual object localizer and 638 volumes for each run of the memory task.

#### fMRI analysis

Most human visual selectivity studies of the ventral temporal cortex have been performed at the group level averaged across participants and oftentimes do not match the functional organization displayed in individual participants. To better understand the precise nature of memory reinstatement in ventral temporal cortex, we implemented the surface-based fMRI analysis at the individual participant level. Preprocessing was performed using fMRIPrep 1.4.0 (Esteban et al., 2019), which is based on Nipype 1.2.0 (Gorgolewski et al., 2011). This pipeline uses a combination of well-known software packages to implement the most suitable tools for each stage of preprocessing. The preprocessed task fMRI data were entered into a general linear model (GLM) to estimate fMRI activity at each vertex/voxel in each run FreeSurfer Functional Analysis Stream (FSFAST) (Burock & Dale, 2000). Linear contrasts were computed to estimate the effects of interest. Fixed-effects analyses were conducted to estimate the average effects across runs within each participant.

#### Functional data analysis

Using the output from the fMRIprep, surface-based vertex-wise GLM was used to analyze the preprocessed MR time series. The GLM included regressors of no experimental interests generated from fMRIPrep preprocessing, including motion estimates. Within-subject first-level analysis was carried out using FreeSurfer FSFAST. For the visual object localizer, the GLM contained 10 regressors-of-interest corresponding to the 10 visual categories: faces, bodies, cars, corridors, instruments, limbs, houses, letters, numbers, and scrambled. For the memory experiment, the GLM contained 5 regressors-of-interest: memory encoding, memory retrieval (hits), memory retrieval (correct rejections), and memory retrieval errors (misses and false alarms).

#### Manual definition of visual selective regions

Face-selective regions on the VTC surface were manually identified for each hemisphere and each participant. First, contiguous sets of cortical surface vertices that passed a common threshold (t>3) of the face-selective map (face > all other visual categories) formed face-selective clusters. Then, these clusters were labeled as mFus, pFus, and IOG based on anatomical landmark markers and criteria outlined in prior work (Chen et al., 2023; Weiner & Grill-Spector, 2010, 2012, 2013). Similarly, for control analyses parahippocampal place area (PPA) was manually identified for each hemisphere and each participant based on the same cluster-forming common threshold (t>3) of the place-selective maps (place>all other visual categories). The clusters were labeled as PPA based on anatomical landmark markers outlined in prior work (Epstein et al., 1999; Kanwisher & Yovel, 2006).

#### Activity-weighted center of mass analysis

To quantify the possible shift in location of neural response from perception to memory encoding and retrieval, we calculated the weighted center of mass for each face-selective area on each surface, where the contribution of each vertex to the center of mass was weighted by its activity map (t-statistic of the contrast) outlined in Steel et al. (2021). In simple cases, the center of mass of an object is usually located at its geometric center, such as the middle of a ball. However, when considering the center of mass for face-selective regions, the distribution of neural activity becomes crucial. Parts of the region with stronger activity will have a more significant impact on the position of the center of mass compared to parts with weaker activity. Consequently, the activity-weighted center of mass reflects where the neural activity is concentrated and is sensitive to changes in the distribution of activity levels. This allows us to investigate whether there is a shift in the anatomical location of activation maps during memory retrieval compared to perception and encoding. Face selectivity centers of mass were calculated using all the vertices within the boundaries of face-selective ROIs identified, namely mFus, pFus and IOG for each hemisphere and each participant, weighted by the t-statistic of the contrast: Face > all other visual categories. Memory encoding activity centers of mass were calculated using the same vertices within the boundaries of ROIs and weighted by the t-statistic of the memory encoding contrast: stimuli presentation > fixation. Memory retrieval activity centers of mass were calculated using the same vertices within the boundaries of ROIs and weighted by the t-statistic of the memory retrieval contrast: hits > correct rejections. The distance between the center of mass for face-selectivity and memory encoding activity and face selectivity and memory retrieval in the y-dimension (anterior-posterior) and x-dimension (medial-lateral) were measured in the unit of millimeters.

### ECoG Methods

#### Patient

Intracranial recordings (electrocorticography, ECoG) were obtained from 1 male patient (20 years of age) undergoing invasive monitoring for the potential surgical treatment of refractory epilepsy at Baylor St. Luke’s Medical Center (Houston, Texas, USA). He provided written and verbal voluntary consent to participate in the experiments reported here. All experimental protocols were approved by the Institution Review Board at Baylor College of Medicine (IRB protocol number H-18112). We excluded electrodes at epileptic foci, and no experiments were recorded in presence of inter-ictal epileptic discharges.

### ECoG experimental design

Taking a similar approach as our fMRI study above, patient performed visual object localizer and word-face memory task (Figure 5 a & b). Details of these tasks are reported above and briefly summarized below.

#### Experiment 1: Visual Object Localizer

In the perception task, the patient was presented grayscale images from 8 visual categories (faces, houses, bodies, limbs, cars, words, numbers, and phase-scrambled noise) in random order (see Figure 5a). On each trial, stimuli were shown for 1000 ms, with a random ISI between 1000 - 1500 ms. During the task the patient were required to press a button whenever they detected a specific stimulus being repeated back-to-back (1-back task). Performance was monitored by an experimenter present in the patient room. A total of 15 different stimuli were presented for each category, with 10 random images being repeated (serving as targets), leading to a total of 160 trials. On average, the task was 7 minutes in duration. The patient performed two runs of this experiment.

#### Experiment 2: Memory task (word-picture paired associates)

In the memory task, the patient performed a paired-associates paradigm with an encoding and retrieval phase. During the encoding phase, the patient was presented with 15 word-face pairs. Each pair was presented for 5000 ms, with a self-paced ISI. The patient pressed a button to advance to the next word-picture pair. After a short delay, the retrieval phase would begin with the patient being presented with only cue words displayed above the box frame, but no picture. Cue words were from a list of 15 old (from the encoding phase) and 15 new words. On each trial of the retrieval phase, the patient was required to retrieve the picture associated with the cue. Cue words were presented for 5000 ms, followed by a response period where patients were asked to provide their memory strength of the cue and associate. On average, the task was 10 minutes in duration. The patient performed three runs of this experiment.

#### Electrophysiological recording

Intracranial EEG data was acquired at a sample rate of 2 kHz and bandpass of 0.3-500 Hz (4th order Butterworth filter) using a BlackRock Cerebus system (BlackRock Microsystems, UT, USA). Initial recordings were referenced to an inverted subdural electrode away from the pathological zones. During recordings, stimulus presentation was tracked using a photodiode sensor (attached to stimulus monitor) synchronously recorded at 30 kHz. All additional data processing was performed offline.

#### Electrode localization and selection

To identify electrodes located within the ventral temporal cortex, a post-operative CT scan was co-registered to a pre-operative T1 anatomical MRI scan for each patient, using FSL and AFNI (Cox, 1996; Dale et al., 1999). The volume location of each electrode was identified by clear hyperintensities on the aligned CT using AFNI and visualized using iELVis software functions in MATLAB (v2016a, MathWorks, MA, USA)(Groppe et al., 2017).

#### Preprocessing and spectral decomposition

All signal processing was performed using custom scripts in MATLAB (v2020b, MathWorks, MA, USA). First, raw EEG signals were inspected for line noise, recording artifacts or interictal epileptic spikes. Electrodes with clear epileptic or artifactual activity were excluded from further analysis. Second, signals were notch filtered (60 Hz and harmonics) and average re-referenced. Finally, Re-referenced signals were down-sampled to 1 kHz and spectrally decomposed using Morlet wavelets, with center frequencies spaced linearly from 2 to 200 Hz in 1 Hz steps (7 cycles).

#### Statistical analysis

fMRI and iEEG data were subject to quantification using parametric and non-parametric statistical methods as appropriate for underlying data distributions. Mixed-effects models, using subject as random effect, were employed to account for individual variability or population sampling nuisance. Confidence intervals of the Spearman correlation were estimated by 1,000,000 replicates of bootstraps and corrected with a bias-corrected accelerated (BCa) method. Statistical analyses were carried out using R statistical software (R Development Core Team, 2010).

## Acknowledgments

This work was supported by NIH grants R01MH116914 to B.L.F and R01EY023336 to D.Y. The authors thank Dr. Kevin Weiner for advising on the characterization of face-selectivity data.

## Author contributions

Y.Y.C. and B.L.F. designed research; Y.Y.C., B.L.F., A.A. and D.Y. performed research; Y.Y.C. analyzed data; Y.Y.C. and B.L.F. wrote the paper.

